# Developing political-ecological theory: The need for Many-Task Computing

**DOI:** 10.1101/871434

**Authors:** Timothy C. Haas

## Abstract

Models of political-ecological systems can inform policies for managing ecosystems that contain endangered species. One way to increase the credibility of these models is to subject them to a rigorous suite of data-based statistical assessments. Doing so involves statistically estimating the model’s parameters, computing confidence intervals for these parameters, determining the model’s prediction error rate, and assessing its sensitivity to parameter misspecification.

Here, these statistical algorithms along with a method for constructing politically feasible policies from a statistically fitted model, are coded as JavaSpaces™ programs that run as compute jobs on either supercomputers or a collection of in-house workstations. Several new algorithms for implementing such jobs in distributed computing environments are described.

This downloadable code is used to compute each job’s output for the management challenge of conserving the East African cheetah (*Acinonyx jubatus*). This case study shows that the proposed suite of statistical tools can be run on a supercomputer to establish the credibility of a managerially-relevant model of a political-ecological system that contains one or more endangered species. This demonstration means that the new standard of credibility that any political-ecological model needs to meet before being used to inform ecosystem management decisions, is the one given herein.

## 1 Introduction

There is a need to acknowledge the complexity of political-ecological systems and the significant challenges to building theories of them [1]. Such systems lie at the interface between social/political science and ecology. The complexity of each of these fields coupled with an additional layer of complexity introduced by the interactions between sociological/political systems and natural systems can result in highly complex system dynamics, i.e., ones that are stiff, nonlinear, and possess feedback loops. For example, Schoon and Van der Leeuw [2] note that systems composed of interacting sociological and ecological subsystems are quick to change and rarely stay in equilibrium for long. Further, many state variables are needed to describe both the decision making processes of the relevant social groups, and the functioning of the involved ecosystem. A *political-ecological system* is also referred to as a *socio-ecological* system or *social-ecological* system (e.g., see [3]). The former term is emphasized herein because those political actions and processes that drive social movements are often initiated by groups seeking to gain increased political power [4]. Building such models is more than an academic exercise. Indeed, the alarming decline in the planet’s biodiversity [5], creates a crucial need for credible political-ecological theory to guide the development of sustainable biodiversity conservation policies. In this article, biodiversity (a shortening of the two words “biological” and “diversity”) is

> an attribute of a site or area that consists of the variety within and among biotic communities, whether influenced by humans or not, at any spatial scale from microsites and habitat patches to the entire biosphere [6].

In addition to the challenge of building political-ecological theory, there is a deeper problem with using such models to guide ecosystem management policy: Unless such a model is shown to be credible using in-part, appropriate statistical methods, any policy recommendations based on output from the model may receive only mixed acceptance by those affected. As argued in [7, p. 181], there is need for a common model credibility standard to be met before the output of a model of a political-ecological system is deemed to be policy-relevant. This is because there may be skepticism towards large scientific models that have not had their parameters statistically estimated nor their parameter sensitivities assessed [8], [9]. These skeptics may be unwilling to cooperate with efforts to implement ecosystem management policies that are based in-part on output from these unassessed models.

But what is a credible model? Patterson and Whelan [10] state that “Model credibility is about the willingness of people to make decisions based on the predictions from the model.” In other words, a model is credible when a decision maker places enough trust in its predictions to use those predictions to select management actions. Call the model’s behavior, functioning, relationships, and systems of equations, its collective *mechanism*. Patterson and Whelan [10] believe the decision maker’s trust is won if (a) the model’s mechanism is based on known principles that govern the phenomenon being modeled; (b) all aspects of the model’s mechanism are testable, i.e., there are observable variables in the model on which data may be collected and used to conduct statistical hypothesis tests of the presence of these behaviors in the real world; and (c) the out-of-sample prediction error of the model’s predictions is below the decision maker’s threshold.

To make the assessment of a political-ecological model’s credibility easier to perform, the present article develops and demonstrates an integrated suite of statistical methods for assessing model credibility components (b) and (c), above. Some of the hypotheses of component (b) may concern the sensitivity of the model to perturbations to its parameters. The testing of such hypotheses is typically referred to as performing a *sensitivity analysis*.

For the remainder of this article, the term “model validation” will not be used because in this author’s opinion, it is too ambiguous a term to support a consensus about whether a valid model can be established at all, let alone how it might be quantitatively assessed (see [11] and [12]).

An *agent-based model* consists of a collection of entities that make a sequence of decisions through time based on their goals and inputs from other agents. As described by Bonabeau [13],

> In agent-based modeling (ABM), a system is modeled as a collection of autonomous decision-making entities called agents. Each agent individually assesses its situation and makes decisions on the basis of a set of rules. Agents may execute various behaviors appropriate for the system they represent – for example, producing, consuming, or selling.

An ABM is often built to model a social system that is too complex to represent using mathematical or statistical models (Bruch and Atwell 2015). In ecology, the word “agent” is often replaced with the word “individual” to emphasize that the entities are individual flora or fauna whose behavior is more genetically defined rather than being based on a belief system such as utility maximization. As the authors of [14] state, *individual-based models* (IBMs) “explicitly represent discrete individuals within an (ecological) population and their individual life cycles.” Grimm and Railsback [15] give a comprehensive treatment of this class of models as used to model natural, nonanthropogenic populations, e.g. trees, insects, plants, fish, or terrestrial mammals. One approach to modeling a political-ecological system is with a combination of an ABM to capture the system’s anthropogenic actions, and an IBM to capture the dynamics of the affected ecosystem. These two submodels interact with each other in order to capture the effects of actions taken by groups of humans that affect the ecosystem – and the feedback effects from the ecosystem back to those groups.

For example, Haas and Ferreira [16] build an economic-ecological model of the rhinoceros (*Ceratotherium simum*) horn trafficking system. This model contains submodels (agents) of rhino horn consumers, rhino poachers, and those antipoaching units attempting to stop the poachers from poaching. These latter two submodels interact with an IBM of the rhino population being illegally harvested. Haas and Ferreira [17] extend the poachers group submodel of this ABM-IBM model by adding a mechanism that explains how these individuals weigh the risk of being prosecuted for poaching against its profit potential. These authors then use this submodel to evaluate the practicality of policies aimed at providing employment opportunities for rhino poachers versus policies that intensify the enforcement of anti-poaching laws. This ABM-IBM model contains several hundred parameters.

### 1.1 Related work

#### 1.1.1 Socio-ecological modelling

In a highly cited article, Macy and Willer [18] discuss how ABMs can advance sociological theory. Conte and Paolucci [19] note the potential that ABMs have for social science theory construction but express concern that current models are delivering over-simplified models of cognitive processes. These authors believe ABMs have the potential to deliver much more cognitively realistic models of their agents. Bruch and Atwell [20] explain how ABMs can help develop policy-relevant social science theory, and then review how to validate such models against sociological data sets.

Within environmental modelling, the authors of [21] build a political-ecological model of land developer agents, homeowner agents, and government agents coupled to a natural model that consists of its own, interacting submodels of land-cover transition, hydrology, and wildlife habitat. Developers seek to develop land parcels, homeowners may decide to protest such development decisions, and government agents work to enforce environmental standards. Another example of a political-ecological model is given in [22]. These authors develop land manager agents who make decisions to buy or sell portions of their land in response to changes in the profitability of the land that, in-turn, is influenced by the land’s species richness, and governmental incentives or rules. A patch-based dynamic model of species presence and absence forms the natural system submodel. Each of these models have at least ten parameters. Neither model is assessed against observations from the real world.

#### 1.1.2 Socio-ecological model parameter estimation

A literature search uncovered only two articles describing the statistical estimation of a socio-ecological model’s parameters, namely, [23], and [17]. Several articles, however, were found on the estimation of either strictly social models or strictly ecological models. A Markov Chain Monte-Carlo (MCMC)-based method is developed in [24] for finding maximum likelihood parameter estimates of a deterministic model of wildlife population dynamics. A three-step method is given in [25] for finding the maximum likelihood parameter estimates of a deterministic model of bacterial population growth. Step 1 consists of translating the differential equation system into a *randomized maximum a-posteriori* (rMAP) form, Step 2 consists of discretizing this function, and Step 3 involves maximizing the likelihood function via an interior point solver. In [26], the parameters of a deterministic model of phytoplankton growth are estimated with least squares and several heuristic optimization algorithms.

There are considerably fewer statistical methods in the literature for estimating the parameters of a stochastic ecosystem model such as the stochastic population dynamics model of a terrestrial species studied in this article. One family of frequentist parameter estimators that can be applied to this problem are *minimum simulated distance estimators* (MSDEs). The word “distance” in MSDE refers to that between two probability distributions, typically one that is strictly data-derived, and one that is generated by a model. This distance can be quantified. One way to do so is to set it equal to the Hellinger distance. For example, in [23], a Hellinger distance-based MSDE is used to estimate the parameters of a stochastic, dynamic model of a political-ecological system. Within biokinetics, Poovathingal and Gunawan [27] use an MSDE to estimate the parameters of a stochastic biochemical model. Within economics, Grazzini and Richiardi [28] use MSDE to fit the parameters of an ABM of stock market traders, and an ABM of consumers adopting a new product. They find their parameter estimates to be minimally biased.

#### 1.1.3 Socio-ecological model sensitivity analysis

A model is sensitive to a set of parameters if small perturbations to their values significantly affects the model’s outputs. Helton and Davis [29] review *probabilistic sensitivity analysis*. The authors of [30] perform a probabilistic sensitivity analysis of a complex salmon population dynamics model. In [31], a probabilistic sensitivity analysis of an agricultural model is performed in order to assess the sensitivity of its output (net present value (NPV)) to misspecified inputs (price, cost, and yield). This author employs *high performance computing* (HPC) to complete the lengthy computations. Based on this experience, this author calls for such HPC to be employed to calibrate model parameters – similar to the statistical estimation of parameter values discussed herein.

#### 1.1.4 Integrated statistical assessment of a socio-ecological model’s credibility

A literature search uncovered no articles describing an integrated statistical assessment of a socio-ecological model’s credibility. One article, however, did give a specific suite of activities to statistically assess an ecosystem model’s credibility. Focusing on linear regression-based forest growth models, Vanclay and Skovsgaard [32] believe the evaluation of an ecosystem model should include (1) an interrogation of the model’s logic to determine whether it is parsimonious and biologically realistic; (2) a statistical estimate of its parameters; (3) point and interval estimates of its prediction accuracy; (4) computation of statistical goodness-of-fit tests; and (5) a probabilistic sensitivity analysis. These authors believe statistical resampling methods have a potential use in their third and fourth recommendations. These authors, however, do not apply their recommendations to a case study, nor implement them in a software package.

Johnson and Omland [33] highlight the distinction between model goodness of fit (GOF) and model selection and note that GOF diagnostics ignore model complexity (number of parameters) and focus exclusively on the model’s fit to data. Yarkoni and Westfall [34] call for a shift in focus from building models that pass in-sample GOF tests towards the building of models that have low prediction error rates (out-of-sample performance). This is particularly true for models that are used to guide decisions aimed at changing the future behavior of a system (out-of-sample). A political-ecological system is, in-part, a model of how humans behave and hence, the focus on prediction for psychological models as advocated by Yarkoni and Westfall applies to political-ecological models. As Yarkoni and Westfall state,”What we will hopefully then be left with are models that can demonstrably do something foundational to the study of psychology: reliably predict human behavior.”

### 1.2 Simulating a political-ecological system

#### Definition 1.1.

A *political-ecological system simulator* (hereafter *simulator*) is an executable computer program capable of approximating the outputs of a stochastic model of a political-ecological system.

Haas [7, p. 5] describes such a stochastic model:

> As a step towards meeting this need, this book describes an Ecosystem Management Tool (EMT) that links political processes and political goals to ecosystem processes and ecosystem health goals. Because of this effort to incorporate the effects of politics on ecosystem management decision making, the EMT described in this book is referred to as a politically realistic EMT or simply the EMT. This tool can help managers identify ecosystem management policies that have a realistic chance of being accepted by all involved groups and that are the most beneficial to the ecosystem. Haas (2001) gives one way of defining the main components, workings, and delivery of an EMT (referred to there as an Ecosystem Management System). The central component of this EMT is a quantitative, stochastic and causal model of the ecosystem being managed and the social groups involved with this management.

In this simulator, influence diagrams (IDs) (see [35, p. 125]) are used to implement sub-models for group decision making, and ecosystem functioning. An ID is a bayesian belief network with deterministic input nodes. For instance, the political-ecological system models of Haas and Ferreira ([16], [36], and [17]) are computationally implemented through their attendant simulators.

The central argument of this article is that for simulators to effectively contribute to the development of political-ecological theory and ecosystem management policies, the following three activities need to be performed in sequence: (1) statistically fitting the simulator’s parameters to data sets of *political-ecological actions* [37], (2) assessing the *credibility* of this fitted simulator, and (3) running computations on this (now) credible simulator to find politically feasible and sustainable ecosystem management policies.

The first of these activities is fundamental to the success of the subsequent two. As an example of the superiority of statistical estimation of a simulator’s parameters relative to other ways of assigning them, the authors of [38] find that an ABM fitted with the statistical method of maximum likelihood estimation produces a model that outperforms the same model calibrated to minimize its root mean squared error (RMSE). Performance is defined therein to be the fitted model’s ability to forecast homeowner adoption of rooftop solar panels.

### 1.3 EMT procedure

The above-mentioned three activities form part of a step-by-step procedure given in [7, pp. 77-78] for using an EMT. A new version of this procedure follows.

**Step 1:** Identify the boundaries of the ecosystem to be managed. Typically, this ecosystem will host one or more endangered species.

**Step 2:** Identify those political groups that directly or indirectly affect this ecosystem. Construct submodels of these groups. Cast these submodels as IDs and express them in the **id** language. This language is part of the **id** software system (see [39]). Use theories of cognitive processing to assign *hypothesis values* to the parameters of these group submodels. Load these values into *hypothesis parameter files* – one file for each group. It is assumed that individuals trained in the cognition of decision making will be involved in constructing these submodels.

**Step 3:** Construct a population dynamics submodel of all species identified in Step 1. Cast this submodel as an ID and express it in the **id** language. Use ecological theory to identify hypothesis values for the parameters of this ecosystem submodel. Load these values into a hypothesis parameter file. It is assumed that individuals trained in ecology will be involved in constructing this submodel.

**Step 4:** Using all of the above files, create a master file that defines the political-ecological system simulator composed of these interacting group submodels and ecosystem sub-model.

**Step 5:** Acquire a data set of political-ecological actions made by some of the groups modeled in Step 2, and the ecosystem modeled in Step 3. The ecological component of this data set might consist of observations on the spatio-temporal abundance of several species.

**Step 6:** Use **id** to statistically fit some subset of the simulator’s parameters to this data set using *consistency analysis*. This statistical estimator (see [23], and [7, pp. 46-52]) delivers parameter estimates that result in the simulator’s probability distributions on its output variables being as similar as possible to empirical distributions derived from data while at the same time being as close as possible to those derived from political-ecological theory.

**Step 7:** Use **id** to compute jackknife confidence intervals for the parameters estimated in Step 6.

**Step 8:** Conduct an analysis of the simulator’s credibility (see [7, pp. 179-198]) by using **id** to perform the two separate jobs of (a) estimating the simulator’s prediction error rate through computation of its one-step-ahead prediction error rates; and (b) performing a *deterministic sensitivity analysis* using thresholds defined by the parameter confidence intervals found in Step 7. If the simulator displays error rates that are no better than blind guessing (all options in each group submodel are equally likely), or it displays unacceptable sensitivity to some of its parameters, re-formulate one or more of the simulator’s submodels and go back to Step 6. Continue in this manner until the simulator is credible.

**Step 9:** Use **id** to run a job with this (now) credible simulator to construct the *most practical ecosystem management plan* (MPEMP) (see [7, pp. 52-53]).

**Step 10:** Implement this MPEMP in the real world.

**Step 11:** As new data becomes available, repeat Steps 6 through 10.

### 1.4 Addressing the computational challenge

Call one execution of the **id** statistical estimation command, a *batch job* or simply, a *job* (see [40], and [41]). In general, let a *simulator job* refer to one execution of the computations needed to either (1) statistically estimate the parameters of a political-ecological system simulator; (2) compute parameter confidence intervals; (3) compute a measure of a simulator’s prediction error rate; (4) perform a deterministic sensitivity analysis; or (5) find, using the simulator, a politically feasible ecosystem management policy. Note that these five simulator jobs are integrated in that the first two jobs share the same estimator, the fourth job needs the confidence intervals found in the second job, and the fifth job uses the fitted model that was found by the first job.

Each of these simulator jobs involves many different algorithms and sub-computations to execute those algorithms. Execution of these sub-computations collectively, results in the job’s final set of outputs. Call each of these sub-computations, a *task*.

Simulator jobs can require large amounts of computer time – orders of magnitude more time than for example, the fitting of a wildlife capture-recapture model with the statistical method of maximum likelihood. The need for large amounts of computer time can become a challenge for those scientists, government agencies, and NGOs needing to run such computations. Hereafter, call these groups and individuals who are involved in biodiversity protection, *ecosystem managers*. The handicap these managers face is that funding to support the active management of ecosystems can be uneven. For example, circa 2017-2019, the United States Environmental Protection Agency (USEPA) is being down-sized by President Trump’s administration [42]. But managing an ecosystem with the goal of conserving its biodiversity requires an on-going analysis of monitoring data as it arrives in real-time in order to guide the development of management actions that, when implemented, result in successful biodiversity outcomes. This means that ecosystem managers need to have alternative computing options should they be temporarily unable to afford supercomputer time from an external HPC provider.

This article argues that a practical way to meet this computational challenge is to implement these jobs as *many-task computing* (MTC) applications. The authors of [43] state that many-task jobs are

> loosely coupled that are communication-intensive but not naturally expressed using standard message passing interface commonly found in high performance computing, drawing attention to the many computations that are heterogeneous but not “happily” parallel.

In other words, jobs that could benefit from distributed computing but, due to their many complex and inter-dependent tasks, existing parallelization tools are difficult to apply. As explained and shown below, JavaSpaces™ technology (see [44]) is a free and easy-to-learn way to program MTC applications that can be run on the computers of an external HPC provider or, if necessary, on a grassroots distributed computing environment formed by a collection of in-house computers.

### 1.5 Article contributions

This article makes three crucial contributions to the development of political-ecological theory and the use of such theory in the formation of politically-feasible ecosystem management policies. These contributions are

1. the first integrated suite of statistical measures for performing parameter estimation and credibility assessment of a political-ecological model and its attendant simulator,
2. a new method for constructing politically feasible and sustainable ecosystem management policies, and
3. downloadable software for implementing these methods as MTC applications via JavaSpaces technology.

## 2 Materials and Methods

First, the statistical theory underpinning each simulator job is given. The Section continues with a review of how a JavaSpaces program can be used to code an MTC application. The Section concludes with algorithms and runtime issues particular to the casting of simulator jobs as MTC applications.

### 2.1 Statistical estimation of simulator parameters

Consistency analysis is a frequentist parameter estimator that is related to MSDE. Hence, Hellinger distance is reviewed first before consistency analysis is described.

#### 2.1.1 Hellinger distance

Following [23, Appendix], and [17, Appendix S3], one way to define the distance between two multivariate probability distributions is as follows. Partition a vector of *p* random variables, **U** into **U**^(*d*)^, and **U**^(*ac*)^ – the vectors of discrete and absolutely continuous random variables, respectively. Absolute continuity can be thought of as a strong version of continuity (see [45, p. 210]). Say there are *d* discrete members of **U**, and *c* continuous members. Hence, *p* ≡ *d* + *c*. Let the *probability density probability function* (PDPF) be

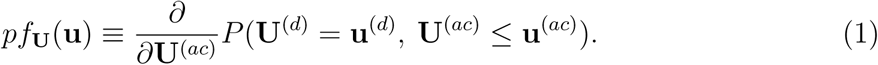

Let **U**|***β*** notate the random vector whose PDPF is parameterized by the components of ***β***. For example, an ID might be composed of *U*_1_ ∼ Bernoulli(*β*_1_) and *U*_2_ ∼ Normal(*β*_2_ + *u*_1_*β*_3_, *β*_4_). The graph of this ID appears in Figure 1, and its parameter vector, ***β*** = (*β*_1_, *β*_2_, *β*_3_, *β*_4_)′.

**Fig. 1.**
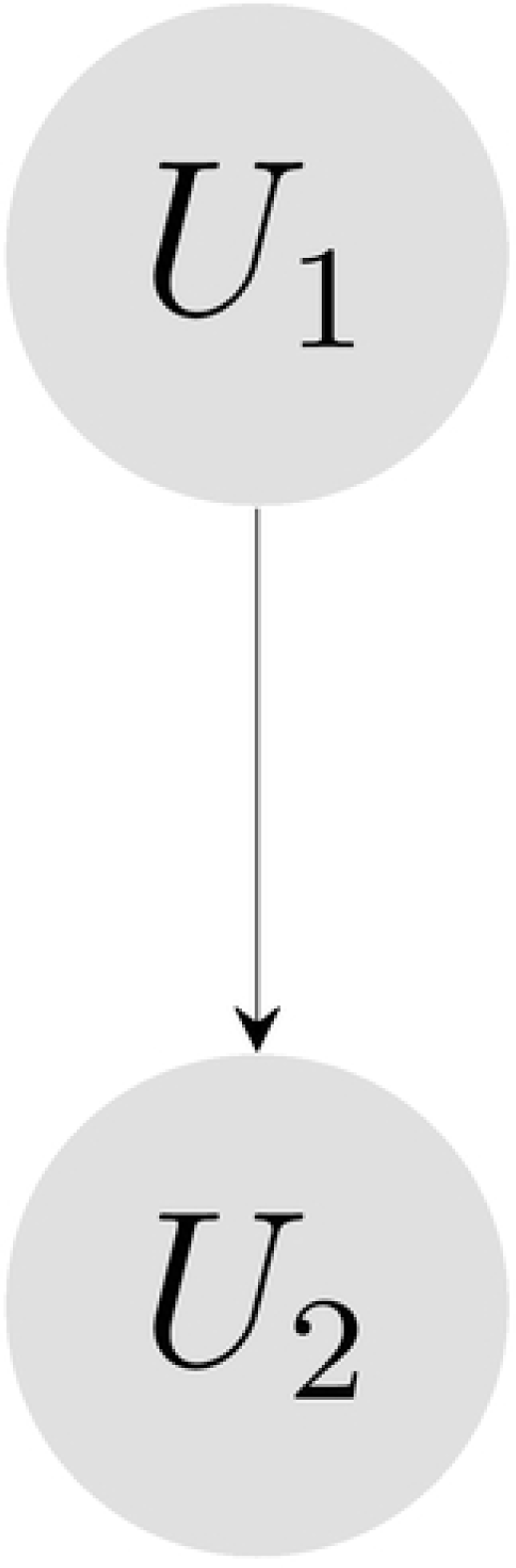
The graph of the ID wherein *U*_1_ influences *U*_2_ and both of these nodes are stochastic (indicated by circles).

In terms of the PDPF, the Hellinger distance between two probability distributions is

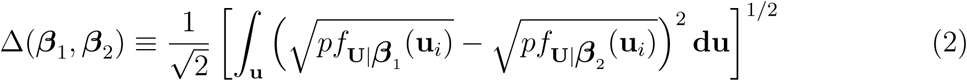

and is bounded between 0 and 1 ([46]).

#### 2.1.2 Consistency analysis

Haas and Ferreira [17] give a description of consistency analysis before applying it to a model the political-ecological system of rhino horn trafficking. An abbreviated version of this description appears here.

##### Definitions

Let *m* be the number of interacting IDs in a political-ecological simulator. Let **U**_*i*_ be the vector that contains all of the chance nodes that make up the *i*^th^ ID (either one of the group submodels or the ecosystem submodel). Let **U**|***β***^(*ij*)^ be the *i*^th^ ID’s multivariate probability distribution parameterized by the entries in ***β***^(*ij*)^ under the *j*^th^ set of conditioning (input) node values. Each parameter in the ID is assigned a point value a-priori that is derived from either expert opinion, subject matter theory, or the results of a previous consistency analysis. Collect all of these hypothesis values into the *hypothesis parameter vector*, 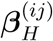. Note that this vector holds the ecosystem manager’s prior beliefs about the point values of the model’s parameters.

Let *l*_*i*_ be the number of belief networks formed by conditioning the *i*^th^ ID on all possible combinations of its input nodes. There are *m* − 1 group submodels, and one ecosystem submodel. Define

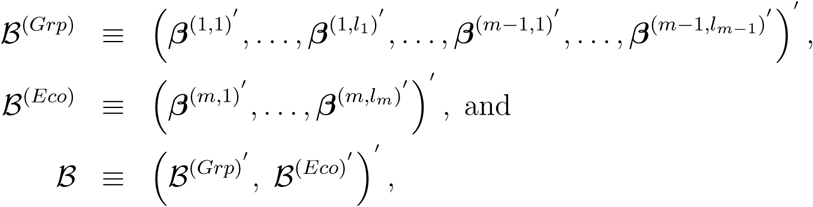

i.e., those parameters that identify all of the group submodels, those that identify the ecosystem submodel, and the collection of all of the model’s parameters, respectively.

As in [7, pp. 17-18], for group submodels, let an *in-combination* be a set of values on the input nodes {*time, input action, actor, subject*}. Let an *out-combination* be a set of values on the input nodes {*output action, target (of that action)*}. A group ID selects an out-combination by computing the expected value of its terminal node, `Overall Goal Attainment` under the received (given) in-combination – and each possible combination of values on the two input nodes of `Out-Action` and `Target`. The out-combination that maximizes this expected value is selected for output.

Let an *in-out pair* consist of an in-combination – out-combination pair. Let *T* be the number of time points at which out-combinations are observed, and 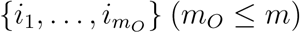 be the set of indices of those group submodels for which at least one out-combination is observed over the observation time interval: [*t*_1_, *t*_*T*_].

Each of the *e* output nodes of the ecosystem submodel is stochastic and corresponds to an observable ecosystem metric. A run of the simulator produces a set of simulated values on each output node at each time point. The mean of these values is an estimate of that node’s expected value at that time point.

Let *g*_*S*_(ℬ) ∈ (0, 1) be a *goodness-of-fit* statistic that measures the agreement of a sequence of out-combinations and/or mean values of ecosystem metrics produced by a simulator and those of a political-ecological actions data set, *S* of observed output actions and/or observations on the ecosystem submodel’s metrics. Larger values of *g*_*S*_(ℬ) indicate better agreement. Let *g*_*H*_ (ℬ) ∈ (0, 1) be a measure of agreement between the multivariate probability distribution on the model’s vector of output nodes that is identified by ℬ, and that identified by ℬ_*H*_. Again, larger values of *g*_*H*_ (ℬ) indicate better agreement. Note that *g*_*S*_(ℬ) is the agreement between a sample and a stochastic model, while *g*_*H*_ (***β***) is the agreement between two stochastic models.

##### Parameter estimator and agreement functions

A consistency analysis is executed with the following four steps.

1. **Specify** the values for ℬ_*H*_.
2. **Initialize** the model’s parameter values by modifying ℬ_*H*_ to form ℬ_*initial*_.
3. **Maximize** the agreement function, *g*_*CA*_(ℬ) (“CA” for “consistency analysis”) by modifying the values of ℬ_*initial*_ to form the vector of *consistent* parameter values, ℬ_*C*_.
4. **Analyze** the differences in parameter values between those in ℬ_*H*_, and those in ℬ_*C*_.

The estimator’s name comes from this final step: analyze the model’s parameters by scrutinizing areas of the subject matter theory that had been used to justify those hypothesis parameter values that, surprisingly, have been found to be very different from their consistent values. This idea of “surprise” is related to the non-bayesian approach to belief revision of *ranking theory* (see [47]). In ranking theory, the model takes the form of a set of propositions and hence, broadly speaking, the value of one of the model’s parameters corresponds to a proposition. These propositions are ranked by the ecosystem manager from completely believable (rank 0) to very unbelievable (rank → ∞), i.e., a very “surprising” proposition. There are several updating rules in ranking theory. These rules do not depend on the size of the data set (the new information), do not require probability distributions on the model’s parameters, and do not involve the calculation of a conditional probability distribution. Belief revision within ranking theory proceeds by computing rank-shifts between the old rankings and the new rankings. These shifts are determined by min{.} operators on these two sets of rankings. The new rankings are assigned based on a subjective interpretation of the new information.

The **Maximize** step of consistency analysis consists of solving

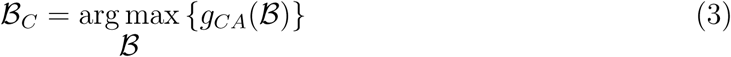

where *g*_*CA*_(ℬ) ≡ (1 − *c*_*H*_)*g*_*S*_(ℬ) + *c*_*H*_ *g*_*H*_ (ℬ), and *c*_*H*_ ∈ (0, 1) is the ecosystem manager’s priority of having the estimated distribution agree with the hypothesis distribution as opposed to agreeing with the empirical (data-derived) distribution. Haas [23, Appendix] gives suggestions for assigning a value to *c*_*H*_. In particular, setting *c*_*H*_ to zero turns consistency analysis into an MSDE. The subjective assignment of *c*_*H*_ in consistency analysis coupled with its role in the solution of (3) is how consistency analysis represents the reliability of the new data – similar to the device used in ranking theory of subjectively re-assigning proposition ranks in the light of new information.

The agreement between the simulator’s hypothesis distributions and the distributions defined by ℬ is 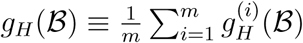 where

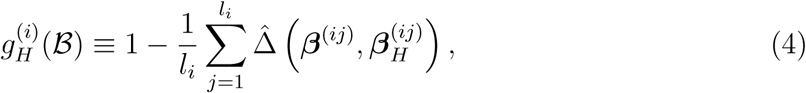

and the estimated Hellinger distance between **U**|***β***_*H*_ and **U**|***β*** is

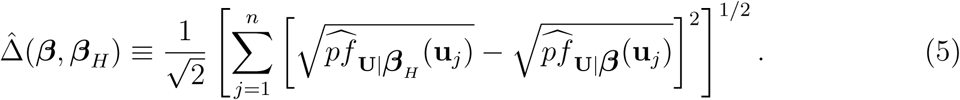

In this estimator, values of the PDPF under an ID’s hypothesis distribution, **U**|***β***_*H*_ and its **U**|***β*** distribution are approximated by first drawing a size-*n* sample of design points from a multivariate uniform distribution on the ID’s chance nodes: **u**_1_, …, **u**_*n*_; and then approximating *pf*_**U**|***β***_(**u**_*i*_) at each of these points with a *k* nearest-neighbor, nonparametric density estimator.

The agreement between observed output actions and those generated by the simulator is

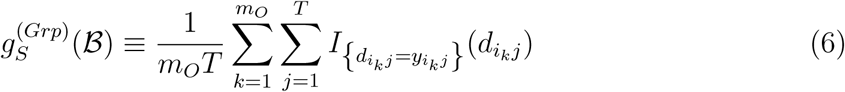

where 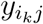 is the observed action of group *i*_*k*_ at time *j*, and 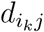 is the submodel-computed action of group *i*_*k*_ at time *j*. Let *S*_*i*_ ≡ {*z*_*i*1_, …, *z*_*iT*_} be the *T* observations on the *i*^th^ ecosystem metric. The agreement between observed outputs of the ecosystem and those generated by the ecosystem submodel is

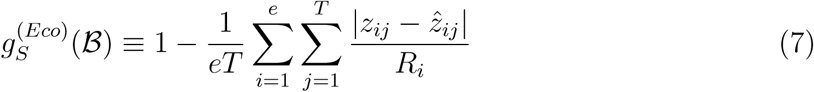

where *R*_*i*_ ≡ max(*S*_*i*_) − min(*S*_*i*_). These latter two agreement functions form the overall data agreement function: 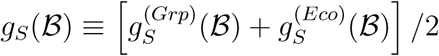.

##### Algorithm for the Initialize step of consistency analysis

The **Initialize** step of consistency analysis is nontrivial due to the discrete nature of the function that counts the number of in-out pairs matched between the data and the simulator’s output. For many different in-combinations, a group may need to select an out-combination that simultaneously maximizes the values of several objectives. The states of these objectives in the group’s present situation is represented in the group’s ID with *situation state* nodes. The perceived states of these objectives upon implementation of a particular out-combination is represented with *scenario state* nodes. Two objectives that are important to several of the groups studied herein are economic objectives, and militaristic objectives. Let situation state, and scenario state nodes take on the values of *negligible (neglig), inadequate (inadeq)*, and *adequate (adequa)*. Also, let a goal node take on the values *unattained (unatta), middling (middli)*, and *attained (attain)*. A group implements a decision option that maximizes the expected value of their `overall goal attainment` (OGA) node.

Based on the decision making theory developed in [7, pp. 83-92], perceived causality in each group submodel is such that situation state nodes are influenced by the input action node; and scenario state nodes are influenced by both the corresponding situation state node and the output action node. In other words, the perceived status of an objective in a scenario is dependent upon its status in the present and the impact of the contemplated output action in the future.

The heuristic: “raise the worst-off objective one level” leads to nine causal sequences (Table 1).

**Table 1.** Patterns of situation state through scenario goal node values used in the **Initialize** step of consistency analysis. SE is *situation economic state*, SM is *situation military state*, SC* is *scenario* * *state*, and SC*G is *scenario* * *goal* where * is either *economic* or *military*. For goal nodes only, let *unatta* correspond to a value of 1.0, *middli* to a value of 2.0, and *attain* to a value of 3.0. The SUM column adds these SCEG and SCMG values to produce an illustrative approximation of the expected value of the OGA node.

| In-Comb. Pattern | SE | SM | Output Action | SCE | SCM | SCEG | SCMG | SUM |
| --- | --- | --- | --- | --- | --- | --- | --- | --- |
| 1 | *inadeq* | *inadeq* | 1 | *neglig* | *inadeq* | *middli* | *unatta* | 3 |
| 2 | *inadeq* | *neglig* | 2 | *neglig* | *neglig* | *middli* | *middli* | 4 |
| 3 | *inadeq* | *adequa* | 3 | *neglig* | *adequa* | *middli* | *attain* | 5 |
| 4 | *neglig* | *inadeq* | 4 | *neglig* | *neglig* | *middli* | *middli* | 4 |
| 5 | *neglig* | *neglig* | 5 | *adequa* | *neglig* | *attain* | *middli* | 5 |
| 6 | *neglig* | *adequa* | 6 | *adequa* | *adequa* | *attain* | *attain* | 6 |
| 7 | *adequa* | *inadeq* | 7 | *adequa* | *neglig* | *attain* | *middli* | 5 |
| 8 | *adequa* | *neglig* | 8 | *adequa* | *adequa* | *attain* | *attain* | 6 |
| 9 | *adequa* | *adequa* | 9 | *adequa* | *adequa* | *attain* | *attain* | 6 |

Haas [7, pp. 166-169] gives an algorithm to initialize the parameters of each group submodel so that the simulator, when run over the time interval of the observed *actions history* (the sample), produces an actions history that matches as many of the observed actions as possible. A new version of this algorithm proceeds as follows.

1. Modify the conditional probability tables (CPTs) of situation state nodes and their parents so that the first nine, most-frequent, different patterns of observed in-combinations (see Table 1) generate nine different patterns of marginal distributions on the economic, and militaristic situation state nodes. Two patterns are different if their modal values are different.
2. Set the CPTs of all scenario state nodes so that the value *inadequate* has the highest value under any combination of the ID’s situation state, and output action nodes.
3. Modify only those CPT entries that carry an output action pattern number given in Table 1 so that they deliver high probabilities on the scenario goal nodes.

Steps 2 and 3 above guarantee that only the output action that is assigned to an in-combination pattern produces a high expected value of the OGA node – and hence enjoys the highest chance of being selected. This algorithm makes no attempt to maintain agreement with the simulator’s set of hypothesis distributions. Such agreement is maximized within constraints during execution of the **Maximize** step of consistency analysis.

The data preparation algorithm forms observed in-out pairs by assuming that a group’s action is a reaction to the immediately-preceding action. This may result in the political-ecological actions data set containing instances where a group is observed to react differently to the same input action on different occasions. Group submodels, however, act as deterministic input-output functions during the execution of the **Initialize** step of consistency analysis. These two characteristics can result in the fraction of matches with the observed in-out pairs being less than one.

### 2.2 Delete-*d* jackknife confidence intervals

The deterministic sensitivity analysis described in the next Section assumes that confidence intervals for each parameter in ℬ are available. One way to find these confidence intervals is to compute *delete-d jackknife confidence intervals* (see [48]). Haas [49, pp. 111-112] gives an algorithm for computing a delete-*d* jackknife confidence interval for a parameter of a stochastic model. This algorithm proceeds as follows.

1. Resample *r* = *n*^0.97^ observations from the observed sample. In other words, temporarily delete *d* ≡ *n* − *r* observations from the observed sample. Politis and Romano [50] show that confidence intervals based on delete-*d* subsamples are consistent if, as *r* → ∞, *r/n* → 0. One way to meet these conditions is to have *r* = *n*^*τ*^ where *τ* ∈ (0, 1).
2. With this *r*-size subsample, compute 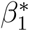, the consistency analysis estimate of the parameter, *β*.
3. Repeat Steps 1 and 2 *n*_*jack*_ times to obtain 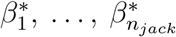.
4. Form a 100(1 − *α*)% confidence interval for *β* by finding the shortest interval that contains (1 − *α*)*n*_*jack*_ of these 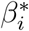 values.

### 2.3 Prediction Error Rates

The simulator’s group submodels produce nominally-valued output in the form of out-combinations. The ecosystem submodel on the other hand, can produce continuously-valued output, e.g. wildlife abundance values. Two different measures of prediction error rate then, are needed. Here, these are the *predicted actions error rate* (*ζ*) for action-target output, and the *root mean squared prediction error rate* (*ϵ*_*i*_) for the *i*^th^ continuously-valued ecosystem metric [7, pp. 186-188].

#### 2.3.1 Predicted actions error rate

Consider a large but finite number of sequential time points, *t*_1_, …, *t*_*T*_. At each of these time points, one or more of the simulator’s group submodels posts one or more out-combinations. Let

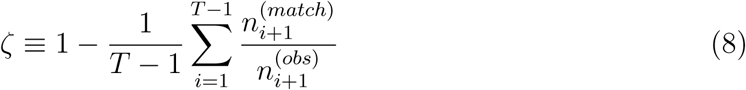

where 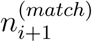 is the number of simulator-predicted out-combinations at time point *t*_*i*+1_ that match observed out-combinations at that time point, and 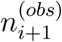 is the number of these observed out-combinations. It is assumed that the simulator’s parameters have been refitted to the political-ecological actions data set using data observed earlier than time point *t*_*i*+1_. The justification for this assumption is that an ecosystem manager would want to refit the simulator as new actions and/or values on ecosystem metrics are observed before using the simulator to predict future group actions and/or future values of ecosystem metrics.

Say that a group submodel has *K* possible out-combinations. In the worst case, one of these out-combinations has a high probability of being chosen at each time point no matter what the input action is. Blind guessing, i.e., assuming all out-combinations are equally likely, would predict this out-combination with probability 1*/K* at each time point resulting in an error rate of about 1 − 1*/K*. An ecosystem manager would prefer the simulator’s predictions over predictions based on blind guessing whenever *ζ* < 1 − 1*/K*.

#### 2.3.2 Root mean squared prediction error rate

Let

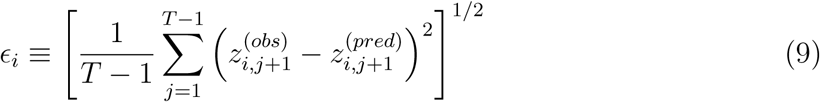

where 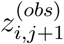 is the observed value of the *i*^th^ continuously-valued ecosystem metric at time point *t*_*j*+1_, and 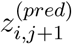 is the simulator’s predicted value of this metric at time point *t*_*j*+1_ where the ecosystem submodel has been fitted to data earlier than time point *t*_*j*+1_. Define an alternative predictor, namely the *naive forecast* to be 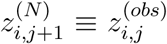 (see [51]). And let *δ*_*i*_ be the RMSE of the naive forecast errors.

#### 2.3.3 Error rate estimation

To estimate these error rates, begin at time point *t*_*s*_, *s* > 0. Then, perform the following two computations at each of the time points 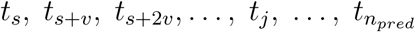 where *v* > 0 is the *refit interval*, 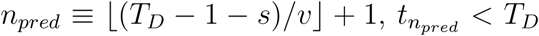, and *T*_*D*_ is the most recent time point in the data set.

1. Re-fit the simulator with consistency analysis using all observed out-combinations up through time *t*_*j*_.
2. Run this refitted simulator from the first time point in the data set up through time point *t*_*j*+1_ to compute predicted values of all output nodes.

With these predictions in-hand, compute an estimate of *ζ* with

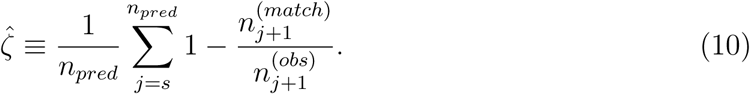

Estimate *ϵ*_*i*_, and *δ*_*i*_ with

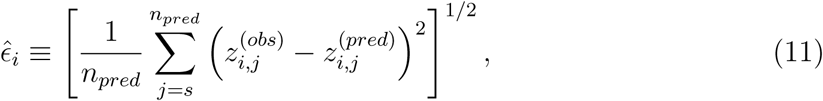

and

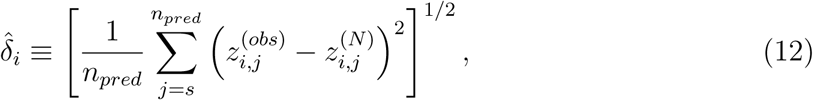

respectively.

Note that the simulator is refitted every *v* time units. Typically, time is measured in years. An ecosystem manager would be constrained by analyst time, computer availability, and data acquisition frequency. A typical refit time interval might be every quarter (three months), i.e., *v* = (4 × 3)*/*52 = 0.2308.

If 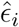 is greater than 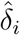, the naive forecast is preferred over the model’s predictions. In this case, the ecosystem manager would be advised to work on refining and/or modifying the model and/or simulator until 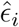 is less than 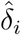.

### 2.4 Deterministic sensitivity analysis

*Deterministic* sensitivity analysis as opposed to probabilistic sensitivity analysis, assesses the sensitivity of a model’s outputs to externally-generated, fixed values of the model’s inputs (see [52]). Haas [7, pp. 182-183] gives an algorithm for studying a simulator’s deterministic sensitivity. A new version of this algorithm is presented next.

#### 2.4.1 Conditions and responses

Input for this algorithm consists of a set of *DSA conditions*, **c**_*DSA*_ (“*DSA*” for “deterministic sensitivity analysis”), and a set of *DSA responses*, **r**_*DSA*_. Each of these sets contains values on simulator submodel output nodes. These values can be those of nominally-valued output action nodes, or those of continuously-valued ecosystem submodel nodes. A particular pair of these sets embodies a counter-example to the types of simulator outputs that the ecosystem manager is hoping to achieve. Typically, a critic or skeptic of the simulator would specify **c**_*DSA*_ and **r**_*DSA*_.

#### 2.4.2 Algorithm

1. Update ℬ_*H*_ to the most recent value of ℬ_*C*_.
2. Specify **c**_*DSA*_, and **r**_*DSA*_ and set the simulator’s time interval accordingly. Place all actions contained in either **c**_*DSA*_ or **r**_*DSA*_ into a file of “observed” actions, and all ecosystem responses contained in **r**_*DSA*_ into a file of “observed” ecosystem outputs. For political actions in either of these sets, initialize ℬ^(*Grp*)^ so that the associated group submodels produce them. And, for any actions in either of these set that are to not happen (referred to here as *complement actions*), initialize ℬ^(*Grp*)^ so that they are not produced by the responsible submodel under any combination of its inputs.
3. Perform the consistency analysis **Maximize** step (see (3)) with this skeptic-postulated actions history (composed of postulated group actions and postulated ecosystem responses). In general, **c**_*DSA*_ and/or **r**_*DSA*_ may contain some mixture of political and/or ecological actions. To ensure a solution is found that results in a close match to all such “observed” group actions and/or ecosystem variables, set *c*_*H*_ to the small value of 0.1 so that the algorithm focuses on matching this skeptic-generated “data” rather than staying true to the hypothesis distributions.
4. Find the parameter in ℬ_*DSA*_ that is the least changed from its value in ℬ_*H*_ relative to its range of scientifically plausible values. Say that it turns out to be the *l*^th^ parameter. Then *β*^(*l*)^ is the most sensitive parameter, and the difference, 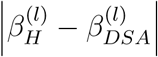 is the accuracy to which this parameter needs to be known. If 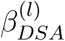 is inside the 95% confidence interval for *β*^(*l*)^ (see Section 1.3, Step 7), or 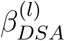 is a scientifically plausible value for *β*^(*l*)^, conclude that this analysis supports skeptic’s concerns about the simulator’s sensitivity to parameter misspecification.

The idea of this algorithm is to search for a set of parameter values that is as close to ℬ_*H*_ as possible but causes the simulator’s outputs to change by an amount that is scientifically significant. If the values in ℬ_*DSA*_ are not statistically different from their consistent counterparts or, are scientifically plausible, then the model’s outputs are *excessively sensitive* to parameter misspecification. This sensitivity in-turn, reduces the credibility of policy recommendations derived from the model’s outputs.

When specifying the condition and response sets with the intention of assessing the sensitivity of group *i*’s submodel, the set **c**_*DSA*_ may contain values on output nodes of submodels other than group *i* while the set **r**_*DSA*_ will be populated exclusively with values on submodel *i*’s output nodes. This is because the simulator may contain submodels whose parameters are sensitive to actions from other groups and/or patterns of ecosystem metric values.

### 2.5 Ecosystem management policymaking

Computing the MPEMP is one way to construct an ecosystem management policy. The algorithm described and demonstrated herein is new. Its development was motivated by earlier algorithms given in [7, pp. 52-53], and [17, Appendix S5]. The idea is to find a set of minimal changes in the beliefs held by ecosystem-affecting groups (relative to their 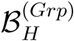 values) so that these groups change their behaviors enough to cause the ecosystem to respond in a desired manner. In other words, the MPEMP is the ecosystem management policy that emerges by finding group submodel parameter values that bring the predicted ecosystem state close to the desired ecosystem state while deviating minimally from 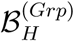.

#### 2.5.1 Definitions

Let **Q**(ℬ) be a random vector composed of a number of the simulator’s ecosystem metrics. For example, **Q**(.) might consist of cheetah abundance, and herbivore abundance in the year 2030. Assume that an ecosystem manager desires the ecosystem to be in a particular state at a particular future time point. This manager expresses this desired state through a set of expected values for **Q**(ℬ). Call this set of desired values, **q**_*d*_. For example, say that it is desired to have 10,000 herbivores and 1,000 cheetah in East Africa in the year 2030. This desired ecosystem state is expressed by specifying

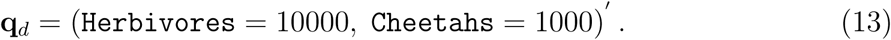

Next, identify those actions that, if taken, would contribute the most towards the ecosystem submodel producing the values in **q**_*d*_. And, identify those actions that, if ceased, would raise the likelihood of the ecosystem submodel producing the values in **q**_*d*_. Collect all of these desirable and undesirable actions into a set called **c**_*MPEMP*_. For example, to achieve these desired values, it is believed that more land should be set aside for wildlife reserves, and poaching should cease. In this case,

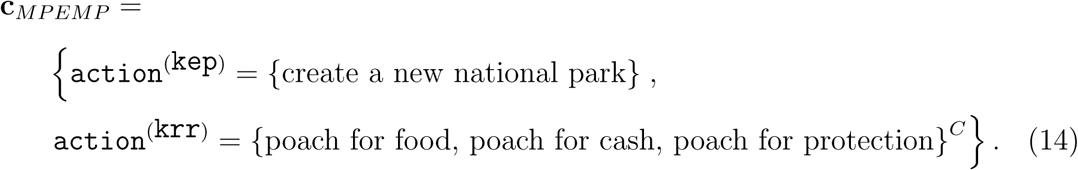

where `kep`, and `krr` are the Kenya environmental protection agency, and Kenya rural residents groups, respectively.

#### 2.5.2 MPEMP algorithm

1. Update ℬ_*H*_ to the most recent ℬ_*C*_.
2. Compute **q**_*H*_ ≡ *E* [**Q**(ℬ_*H*_)].
3. Specify **q**_*d*_ and **c**_*MPEMP*_.
4. Compute initial values for ℬ(*Grp*) with the **Initialize** algorithm of consistency analysis (see Section 2.2.3).
5. Compute

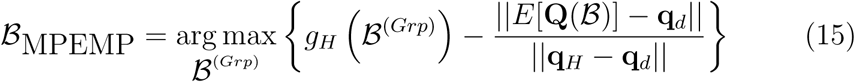

under the set of constraints specified by **c**_*MPEMP*_.

Note that during the search in Step 5, 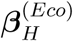 is unchanged. This algorithm implements one way to quantify the concept of a practical ecosystem management policy: associate political feasibility with the value of 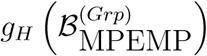 where 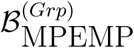 contains the parameters of the decision making submodels whose values have been modified from those in 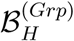 in such a way that now, the sequence of output actions taken by the different groups in the simulator cause a desired ecosystem state at a designated future time point.

A measure of a plan’s political practicality can be defined as

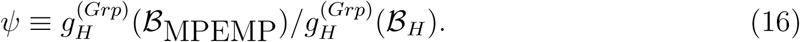

A plan having a value of *ψ* close to 0.0 will face significant political resistance to its implementation because significant changes to the belief systems of one or more groups needs to happen, while one with a value close to 1.0 should not face such stiff resistance.

### 2.6 Coding simulator jobs as MTC applications

The five simulator jobs described above can be computationally expensive. These jobs, however, are not easily organized into parallel, independent tasks but rather, can only be partially parallelized by breaking each of them into sets of dependent tasks that engage in various amounts of data transfer between themselves. For example, a complex task such as function optimization is not easily programmed to run on *graphics processing units* (GPUs) that can process only independent sequences of pipelined floating point operations. Such a set of complex, inter-dependent tasks fits the definition of an MTC application.

But what is the most efficient and cost-effective way to execute MTC applications? One way is to run them on cluster computers. This option is motivated by the work of Raicu and his coworkers [43] who find that an MTC application can be efficiently run on a cluster computer. The authors of [53] assess how efficiently MTC applications run on other computer architectures. A cluster computer consists of a large number of so-called personal computers (PCs) that are connected to each other through high speed interconnects. It is run by an operating system that can assign tasks to one or more of these PCs. An individual PC in the cluster is called a *compute node*. A compute node may possess multiple processors (also known as *cores*). Cluster computing is the dominant architecture of HPC machines. For example, the authors of [54] study Big Data analytics on HPC architectures. All of the architectures considered therein are cluster-based.

As Raicu and his coworkers [43] note, there are many advantages to running MTC applications on a cluster computer as opposed to running in the Cloud or on a heterogeneous collection of PCs. These include

1. I/O systems on cluster computers can be much faster than on other hardware configurations.
2. A core-hour on a cluster computer is often less expensive than many alternatives.
3. Cloud systems and heterogeneous collections of PCs are typically not as reliable as cluster computers.
4. Cluster computers are often fast enough to produce results in a useful period of time.

In order to actually run the five simulator jobs, computer programs need to be written, compiled, and executed on computer hardware. Translating the mathematical expressions of Sections 2.1-2.5 into a programming language is performed by writing code within an *application program interface* (API) that is designed to support the development of task-based parallel programs. A *runtime system* is invoked to execute such programs on computer hardware. This runtime system delivers data and instructions to individual compute nodes, starts these compute nodes, collects output and delivers it to a pre-programmed recipient. The runtime system also detects faults on compute nodes and within compute node processes and delivers the consequential fault information to preprogrammed recipients. Indeed, the action of starting a job on a compute node is a small part of the suite of inter-connected instructions and events that is needed to execute an MTC application.

Many programs written for cluster computers use the *message passing interface* (MPI) (see [55]) to communicate between compute nodes. But, as Dursi [56] notes, MPI is a 25 year old API and, unfortunately for modern MTC applications, operates at too low of a level. This is because its basic abstraction level is that of a message and hence remains “…essentially at the transport layer, with sends and receives and gets and puts operating on strings of data of uniform types.” Dursi [56] reviews the consequences of this low level of abstraction:

> Programming at the transport layer, where every exchange of data has to be implemented with lovingly hand-crafted sends and receives or gets and puts, is an incredibly awkward fit for numerical application developers, who want to think in terms of distributed arrays, data frames, trees, or hash tables. Instead, with MPI, the researcher/developer needs to manually decompose these common data structures across processors, and every update of the data structure needs to be recast into a flurry of messages, synchronizations, and data exchange. And heaven forbid the developer thinks of a new, better way of decomposing the data in parallel once the program is already written. Because in that case, since a new decomposition changes which processors have to communicate and what data they have to send, every relevant line of MPI code needs to be completely rewritten. This does more than simply slow down development; the huge costs of restructuring parallel software puts up a huge barrier to improvement once a code is mostly working.

For complex computations such as the ones described herein, a higher level of abstraction is needed such as that of a task. The authors of [57] review APIs and runtime systems that are designed to support MTC applications. These authors refer to a particular combination of an API and a runtime system as a *task-based parallelism technology* and note that HPC is moving away from the message passing paradigm to such technologies. In order to illustrate why such a move is needed to make progress in HPC, Dursi [58] gives a detailed comparison between MPI programs and those written in other, more modern task-based parallelism technologies.

As identified in [57], an ideal API would have the ability to partition, synchronize, and cancel tasks; specify compute nodes for workers to run on; start/stop workers; receive task or process fault information; and checkpoint a job should a nonrecoverable fault occur. These authors also believe that an ideal runtime system would automatically distribute data and code to workers; schedule workers; and deliver fault information to the master compute node. In addition, the present author believes that in order to bring many-task computing within reach of ecosystem managers possessing only minimal programming skill, the API needs to be easy-to-learn, and use operators whose syntax and semantics are independent of specific runtime systems and computer hardware configurations.

#### 2.6.1 JavaSpaces programs

One way to implement an MTC application is through the JavaSpaces task-based parallelism technology [59]. A JavaSpaces program can support the *master-worker architecture* wherein a master program runs on one compute node having a unique Internet Protocol (IP) address along with *n*_*W*_ workers who run on other, internet-accessible compute nodes and busy themselves by executing tasks that have been posted by the master on a JavaSpace bulletin board. An application is solved via the *bag of tasks* model wherein tasks are distributed by the master across available workers. The master does this by posting tasks on a space, and collecting completed tasks from that space. Batheja and Parashar [60] note that

> This approach supports coarse-grained applications that can be partitioned into relatively independent tasks. It offers two key advantages: (1) The model is naturally load-balanced. Load distribution in this model is worker driven. As long as there is work to be done, and the worker is available to do work, it can keep busy. (2) The model is naturally scalable. Since the tasks are relatively independent, as long as there are a sufficient number of tasks, adding workers improves performance.

And, Noble and Zlateva [61] find that “The simplicity and clean semantics of tuplespaces allow natural expressions of problems awkward or difficult to parallelize in other models [62].” Further, Batheja and Parashar [60] address the runtime system component of JavaSpaces:

> A JavaSpace program provides associative lookup of persistent objects. It also addresses fault-tolerance and data integrity through transactions. All access operations to objects in the space such as read/write/take can be executed within a transaction. In event of a partial failure, the transaction either completes successfully or does not execute at all. Using a JavaSpaces-based implementation allows transacting executable content across the network. The local instances of the Java objects retrieved from the space are active, i.e. their methods can be invoked and attributes modified. JavaSpaces provides mechanisms for decoupling the semantics of distributed computing from the semantics of the problem domain. This separation of concerns allows the two elements to be managed and developed independently [19]. For example, the application designer does not have to worry about issues such as multithreaded server implementation, low level synchronization, or network communication protocols.

In sum, the advantages of a JavaSpaces task-based parallelism technology are:

1. A high level of abstraction: The future of computing lies with clusters of cluster computers. These computing environments will be fully utilized when scientists can write programs that can call other large programs without regard as to how these other programs perform their tasks.
2. Asynchronous, high-level coordination of simultaneous tasks.
3. Communication protocol is outside of the application code so that scientists need not spend time learning and programming inter-processor communications.
4. Internet-aware: Tasks may be executed by any worker that is reachable through a Universal Resource Locator (URL).
5. Fault-tolerant: Dursi [56] shows that processor failure is almost certain during a job that employs thousands of processors. The authors of [63] and [59] both argue that this feature makes JavaSpaces a very attractive tool for HPC applications.
6. Scalable: Only one code need be written and maintained to run jobs on hardware ranging from laptop computers to cluster computers. This natural adaptability of JavaSpaces programs to heterogenous computing platforms was recognized shortly after JavaSpaces was announced [64]. These authors also note that an additional advantage of JavaSpaces is that its learning curve is not high – and that this advantage is often overlooked in evaluation exercises that are solely focused on runtime performance.

Gigaspaces is a particularly simple and efficient implementation of JavaSpaces technology. Specifically, the authors of [65] find that Gigaspaces programs exhibit less inter-compute node communication latency than do JavaSpaces programs executed within other runtime systems. The primary operations on a Gigspaces space are `write`, `read`, `change`, `take`, and `aggregation` [66], [67]. Note that although a JavaSpaces program can support communication between specific workers independent of the master [44, pp. 108-116], such a program would not have high fault tolerance because the recipient compute node of such a message may become unavailable just after the message is sent.

In summary, the JavaSpaces task-based parallelism technology is much more than simply a way to start a Java program. Rather, it is an inter-task communication protocol that is asynchronous and anonymous. A JavaSpaces program starts workers, collects worker outputs, adjusts for faults, partitions tasks, and synthesizes the results of completed tasks. All of these activities can be programmed without the need to learn a language for the micro-management of memory and/or task execution. S1 Appendix A contains shell scripts that start and run a JavaSpaces program on a cluster computer. And S1 Appendix B contains practical guidance for running a JavaSpaces program on a shared cluster computer.

##### Optimization with JavaSpaces

Optimization of stochastic functions under nonlinear constraints can be implemented in a JavaSpaces program via the *multiple dimensions ahead search* (MDAS) algorithm of Haas [7, pp. 219-225]. This algorithm is a parallel version of a nonlinear, constrained optimization algorithm, namely the classic Hooke and Jeeves coordinate search algorithm [68].

MDAS executes by having the master assign each worker a vector of parameter values with which to compute the value of the objective function. These vectors are chosen such that the next *M* parameters are searched simultaneously for a minimum. Each worker computes the objective function value at its assigned set of parameter values. Once all of the workers have returned their function evaluation values to the master, the master checks these values for a new minimum (called an *improvement*). If found, the master stores this new best solution. This parallel search is repeated on these dimensions until no improvements are found. Then, the algorithm moves on to the next *M* dimensions. For *M* = 1, MDAS is equivalent to the (sequential) Hooke and Jeeves algorithm.

Running MDAS with *n*_*W*_ = 8 workers (*M* = 2) gives worst-case, a four-times speedup of an optimization job relative to running the algorithm with only one worker. This is because the sequential version’s inner `for` loop may need to perform up to 2*K* function evaluations before an improvement is found. For *M* = 3, MDAS amounts to an evaluation of all possible visited locations for the next three dimensions in the inner `for` loop of the sequential version. This requires 2 + (3 × 2) + (3 × 3 × 2) = 3^3^ − 1 = 26 parallel evaluations of the objective function. When there are at least *n*_*W*_ = 26 workers available to perform these tasks in parallel, MDAS delivers a six times speed-up over the worst-case of sequential Hooke and Jeeves search when *K*, the number of parameters to be fitted, is a multiple of three. In general, to produce a 2*M* speed up over worst-case sequential Hooke and Jeeves, MDAS needs to be run on a cluster computer having *n*_*W*_ = 3^*M*^ − 1 workers. For example, running with *n*_*W*_ = 242 workers (*M* = 4) gives worst-case, an order of magnitude speedup – and to achieve a 20-fold worst-case speedup (*M* = 5), *n*_*W*_ = 59048 workers are needed. As these speedup values suggest, a guaranteed way to speedup MDAS is by increasing the number of compute nodes that the optimization job can access. Put another way, the inefficient use of a geometrically increasing number of workers is traded for guaranteed worst-case reductions in runtime.

The MDAS algorithm requires master-worker communication at every step (through the collection of results, identifying the new best-solution, and posting of new points at which to evaluate the objective function). Therefore, MDAS is not an *embarrassingly parallel* algorithm. An embarrassingly parallel job (in the sense of “an embarrassment of riches,” see [69]) consists of a set of tasks that can be executed in parallel with no inter-task communication. Also, the objective function evaluation tasks are complex involving for example, the running of a political-ecological simulator many times to support the computation of the consistency analysis objective function. This complexity is qualitatively higher than sending messages to update particular memory locations as is typical in an MPI-based parallel program.

There are of course, other algorithms for performing function optimization on a cluster computer. MDAS is used here, however, because its worst-case speedup characteristics are known; it is scalable; and, because it only requires solution vectors to be sent out to workers but not sent back, it has reduced inter-compute node communication overhead relative to other parallel optimization algorithms such as the simulated annealing-based algorithms developed in [70] and [71]. Further, unlike algorithms such as simulated annealing, it always makes small steps from a feasible starting point and hence is less prone to becoming trapped in an infeasible region. This latter property is crucial when working with a function that has a complicated feasible region boundary. Here, such boundaries typically arise during the optimization of (a) the consistency analysis objective function, (b) the deterministic sensitivity analysis objective function, or (c) the MPEMP objective function.

#### 2.6.2 Simulator job-specific algorithms and runtime issues

Algorithmic details for how each simulator job is converted to an MTC application follow.

##### Consistency analysis

Consistency analysis is run as an MTC application on a cluster computer by performing its **Maximize** step with the MDAS algorithm wherein each worker runs on its own compute node. This makes consistency analysis a straightforward MTC application as it requires simply one cluster computer running one JavaSpaces program. In order to both speedup evaluation of the objective function and to improve the optimization run’s convergence behavior, smooth objective functions are employed in-lieu of those based on the approximate negative Hellinger distance for 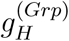, and 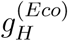 (see (4)). These functions are the negative of the Euclidean distance between the parameters at their hypothesis values and those at a particular trial point in the optimization run. Call these Euclidean agreement measures 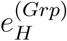, and 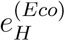, respectively. Although maximizing these Euclidean agreement measures does not guarantee that a point will be found that solves (3), experience described next suggests that a point close to this maximal point can indeed be found.

##### Jackknifing

Jackknifing involves executing consistency analysis on each of *n*_*jack*_ separate delete-*d* sub-samples. It can be implemented as an MTC application by performing all of these *n*_*jack*_ consistency analysis tasks simultaneously. These *n*_*jack*_ consistency analysis tasks are independent of each other and hence may be computed in parallel with no inter-task communication, i.e., this algorithm is embarrassingly parallel. Call this set of tasks the job’s *outer loop*. Nevertheless, the computational expense is high as now, the *n*_*jack*_ consistency analysis tasks require *n*_*jack*_(3^*M*^ − 1) workers.

Running simultaneous optimization tasks is accomplished by running *n*_*jack*_ separate MDAS algorithms in parallel. This is done by adding an inner loop to the MDAS algorithm so that for a given set of *M* dimensions, the objective function is independently evaluated for each jackknife subsample at each solution point that is called for at this set of dimensions.

##### Prediction error rate

Converting this simulator job to an MTC application involves running a consistency analysis task on each of *n*_*pred*_ subsamples (see Section 2.3.1). This is accomplished the same way that the jackknife subsamples are processed.

##### Deterministic sensitivity analysis

The computational demands of a deterministic sensitivity analysis accrue from the consistency analysis performed in its Step 3 (see Section 2.1.2). See above for how consistency analysis is implemented as an MTC application.

##### MPEMP computation

The computational demands of an MPEMP simulator job accrue from the optimization problem solved in the MPEMP algorithm’s Step 5 (see Section 2.5.2). Hence, as with consistency analysis, an MPEMP job is implemented as an MTC application by performing this optimization with MDAS wherein each worker runs on its own compute node.

### 2.7 Case study description

The following Results section contains a case study that applies the five simulator jobs to the credibility assessment and MPEMP computation of an EMT for the conservation of cheetah in East Africa. All input files for this simulator are available at [72]. Hereafter, this simulator is referred to as the *cheetah EMT simulator*.

#### 2.7.1 Overview of the Cheetah EMT simulator

Haas [7, pp. 97-121] builds a simulator of the interactions between cheetah and humans in the East African countries of Kenya, Tanzania, and Uganda. The model consists of group submodels for each country’s presidential office (`kpr`, `tpr`, `upr`), environmental/wildlife protection agency (`kep`, `tep`, `uep`), non-pastoralist, rural residents (`krr`, `trr`, `urr`), and pastoralists (`kpa`, `tpa`, `upa`). In addition, a submodel is built to represent the group of conservation NGOs who have operations in at least one of these countries (`ngo`). All of these group submodels can interact with each other. And, each country’s environmental protection agency, rural residents, and pastoralists submodels can directly interact with a submodel of the ecosystem that spans these three countries (`ecosys`). This ecosystem hosts populations of cheetah and their herbivore prey. This model is formally documented in S1 Appendix C.

An automatic data acquisition system has been gathering data since January, 2007 on this political-ecological system (see [37]). This data set contains 1555 actions observed from the year 2002 to 2019. S2 Data contains this data set. A portion of this data reveals a complex pattern of group actions followed by reactions from other groups (Figure 2). Cheetah abundance data is taken from [73], [74], and [75].

**Fig. 2.**
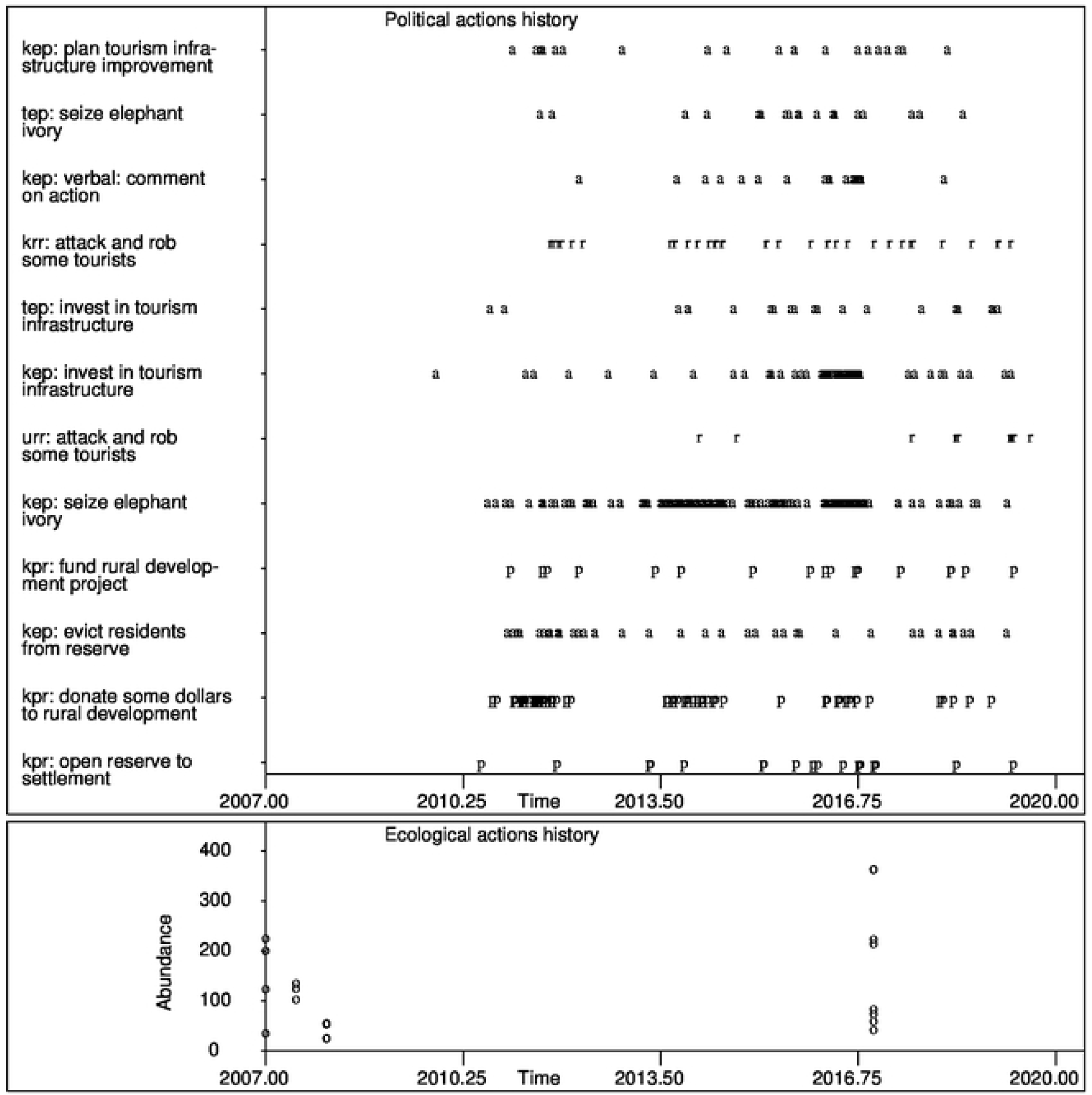
Observed actions history from East African online news stories for the period from January 2007 through June 2019. The symbol “p” indicates an action taken by a presidential office, “a” an action taken by an EPA, “r” an action taken by rural residents, “s” an action taken by pastoralists, and “n” an action taken by an NGO. Selected out-combinations only are labeled. The bottom plot is observed cheetah abundance.

## 3 Results

### 3.1 Consistency analysis

Consistency analysis was used to estimate the parameters of the node: `scenario imminent interaction with police` within the Kenyan rural residents group submodel. A time step of 13 days results in each time interval containing about five actions. The **Initialize** step of consistency analysis (see Section 2.2.3) was run to produce a set of initial parameter values. For this run, each belief network was simulated with 2000 Monte Carlo realizations. Finding the best set of in-out pairs required 4.74 hours on a single PC.

The initial match fraction (the ratio of the number of observed actions matched by the simulator’s output to the number of observed actions) is 0.646. The fraction of actions matched regardless of whether the target was matched, is 0.772, and the corresponding target match fraction is 0.870. See Table 2 for individual submodel match fractions.

**Table 2.**
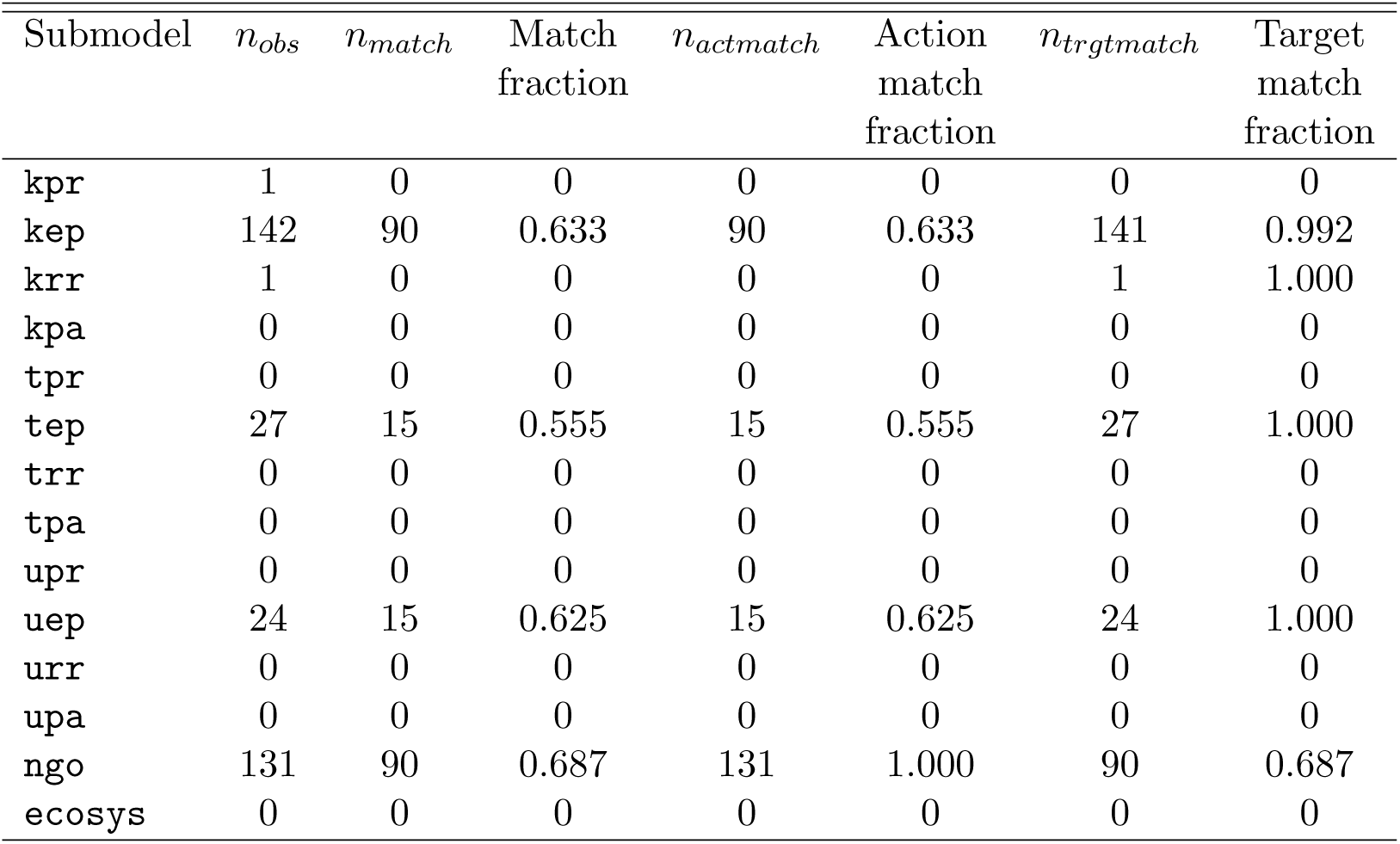
Match fractions from the **Initialize** step of consistency analysis for the cheetah EMT simulator.

Next, the **Maximize** step of consistency analysis was run on the Triton Shared Computing Cluster (TSCC) at the San Diego Supercomputer Center [76]. For this run, *c*_*H*_ was set to 0.99, and each belief network was simulated with 1000 Monte Carlo realizations. Nine compute nodes were employed and the maximum number of function evaluations was set to 1200. Only those parameters having an initial value different from their hypothesis value were modified. This resulted in only 40 of the 459 parameters being active during the optimization run – a significant reduction in the problem’s dimensionality. Initial and final values under the stochastic agreement measure for *g*_*H*_(.) (4) were computed using 5000 Monte Carlo realizations for each belief network.

Under this configuration, the simulator job’s wall clock time was 4.42 hours. The solution achieved a 25.5% increase in *g*_*CA*_(ℬ) (Table 3). Further, the device of maximizing a Euclidean distance-based measure of agreement between the hypothesis and consistent probability distributions did indeed result in an increase in the Hellinger distance-based measure of agreement (Table 3).

**Table 3.** Consistency analysis agreement measures for the cheetah EMT simulator.

| Agreement Measure | Initial Value | Final Value |
| --- | --- | --- |
| $g_S^{(Grp)}(\mathcal{B})$ | 0.6308 | 0.6000 |
| $e_H^{(Grp)}(\mathcal{B})$ | -41.6800 | -29.4314 |
| $g_H^{(Grp)}(\mathcal{B})$ | 0.8468 | 0.8888 |
| $g_{CA}(\mathcal{B})$ | -1.1394 | -0.8483 |

### 3.2 Delete-d jackknife confidence intervals

Jackknife confidence intervals were computed for the parameters that define the `scenario imminent interaction with police` node in the Kenya rural residents submodel of the cheetah EMT simulator. The jackknife subsample size is *r* = 546^0.97^ = 451, and *n*_*jack*_ = 5. These five subsamples were used to compute 50% confidence intervals. Nine compute nodes ran for 4.85 wall clock hours to complete the job. All parameters are significantly different than zero. The five widest confidence intervals (Table 4) indicate that estimates of the group’s beliefs about being prosecuted for actions they might take are not excessively affected by sampling variability.

**Table 4.** The five widest confidence intervals of parameters defining the node `scenario imminent interaction with police` in the Kenya rural residents submodel. These parameters are conditional probability values and hence take values on the unit interval.

| Parameter | Lower Boundary | Upper Boundary | Width |
| --- | --- | --- | --- |
| 168 | 0.110 | 0.362 | 0.252 |
| 165 | 0.161 | 0.412 | 0.251 |
| 171 | 0.211 | 0.462 | 0.251 |
| 174 | 0.111 | 0.262 | 0.151 |
| 183 | 0.111 | 0.262 | 0.151 |

### 3.3 Prediction error rates

Prediction error rate was estimated by computing one-step-ahead predictions of actions, and cheetah abundance from 2016.9 through 2018. This run required 3.25 wall clock hours on the TSCC running nine compute nodes. The run produced 57 predictions resulting in 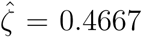, and 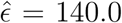 for the cheetah abundance metric. The simulator was refitted to data five times.

### 3.4 Deterministic sensitivity analysis

Say that the ecosystem manager wishes to use the simulator’s outputs to justify his/her position that reducing poaching would slow or reverse the decline in cheetah abundance. A skeptic, however, believes that scientifically plausible parameter values in the cheetah submodel can be found such that when the model is run from 2019 through 2025 under the restriction of no poaching actions, cheetah abundance in the year 2025 will be insignificantly different than that produced by the simulator when run under the assumption that current poaching rates continue into the future. If such parameter values can be found, the skeptic would argue that the model is unable to inform management action selection because the model can be calibrated to either recommend increased antipoaching effort or not recommend increased antipoaching effort.

To represent this skeptic’s belief, **c**_*DSA*_ consists of the single constraint: *no poaching actions occur from the present through the year 2025*, i.e.,

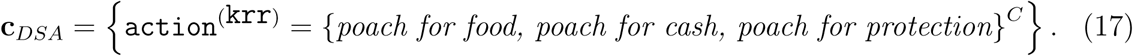

And, **r**_*DSA*_ is populated with predictions of expected cheetah abundance in the year 2025 across several regions in Kenya (Table 5). These predicted values are found by running the simulator out to the year 2025 under the consistent parameter values found in Section 6.2. It is the use of these consistent values that forces poaching rates from 2019 through 2025 to be equal to current poaching rates.

**Table 5.** Cheetah abundance predictions in five regions of Kenya for the year 2025 computed under consistent parameter values. These values make up the set **r**_*DSA*_.

| Region | Abundance |
| --- | --- |
| Laikipia | 200 |
| Samburu | 200 |
| Tsavo | 145 |
| Marsabit | 200 |
| Turkana | 40 |

The mathematical programming problem (3) with variables consisting of the ecosystem submodel’s parameters was solved over the interval 2019 through 2025 and required one hour of wall clock time on the TSCC utilizing eight worker nodes. Initial parameter values were set to ℬ_*H*_ with the exception that values in ***β***^(krr)^ were adjusted as necessary so that any contemplated poaching action produced a small value of *E*[*OGA*]. Doing so caused the Kenya rural residents group to avoid poaching actions during the optimization.

If a solution to (3) were found such that all values in ℬ_*DSA*_ were scientifically plausible, then the skeptic’s position would be supported. As Table 6 indicates, however, the skeptic’s position is not supported because the value for the initial death rate, *r*_0_ (see S1 Appendix C) needed to respect the conditions in **c**_*DSA*_ and the responses in **r**_*DSA*_, is unrealistically high (0.510) under minor poaching pressure.

**Table 6.**
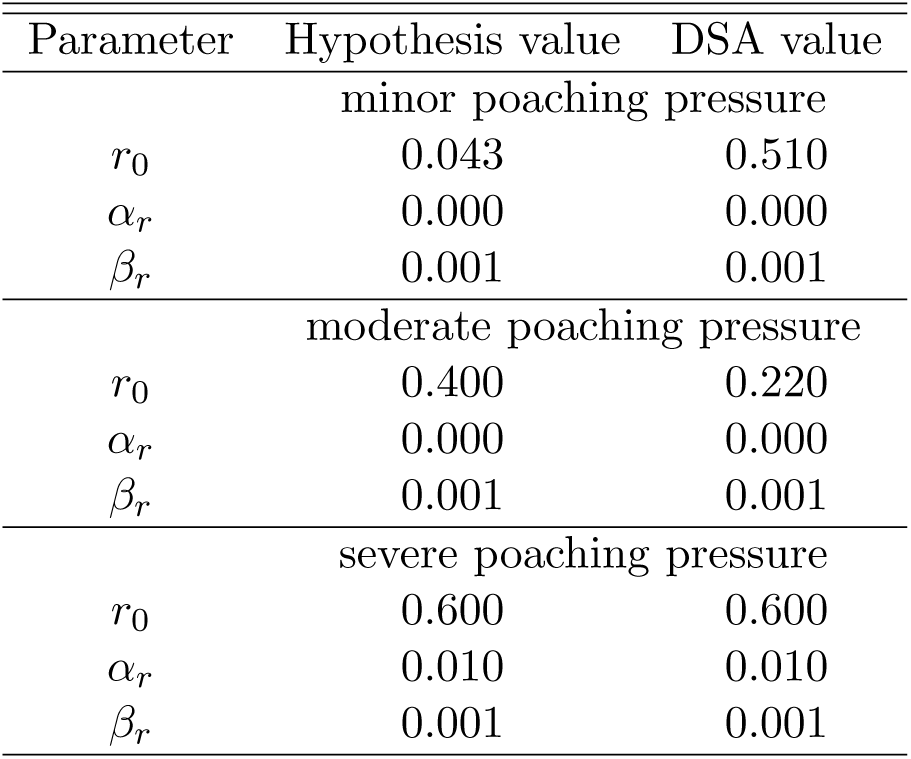
Results for the deterministic sensitivity analysis of the ecosystem submodel.

### 3.5 Overall credibility assessment of the Cheetah EMT simulator

The cheetah EMT model’s mechanism reflects principles of how political-ecological systems function [7, chs. 6-8]. Hence, component (a) of the Patterson and Whelan [10] criteria (see Section 1) is satisfied. Statistical estimation of the model’s parameters is the foundational step for establishing components (b) and (c). The model’s confidence intervals indicate that a selection of the model’s parameters cannot be ignored and can be estimated without excessive uncertainty. The model’s prediction error rates, however, are high. Finally, the model is resistant to a skeptic-created scenario engineered to show the model being unable to inform management action selection.

### 3.6 Finding the MPEMP

Say that it is desired to have 5,000 herbivores and 500 cheetah in East Africa in the year 2030. These target values are expressed by specifying

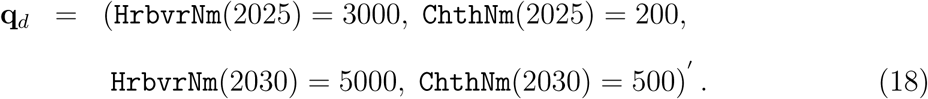

To achieve this ecosystem state, more land needs to be set aside for wildlife reserves, and poaching needs to cease. These conditions are expressed by setting

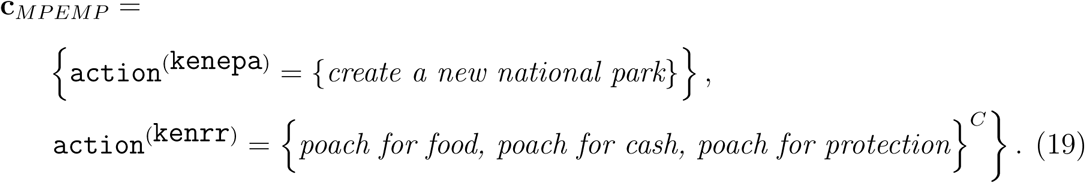

Group beliefs that are to be changed are those of the `imminent interaction with police` node of the Kenya rural resident group.

The simulator job for finding the MPEMP formed a 108-dimensional optimization problem. When run with eight worker nodes on the TSCC, this simulator job required 2.97 wall clock hours to complete. Initial and final values of 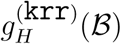 (4) were computed using 5,000 Monte Carlo realizations for each belief network. The MPEMP actions history (Figure 3) is such that Kenyan rural residents substitute the action *verbally protest national park boundaries* for poaching actions. In spite of this behavioral change, however, cheetah abundance does not attain the desired level by the year 2030.

**Fig. 3.**
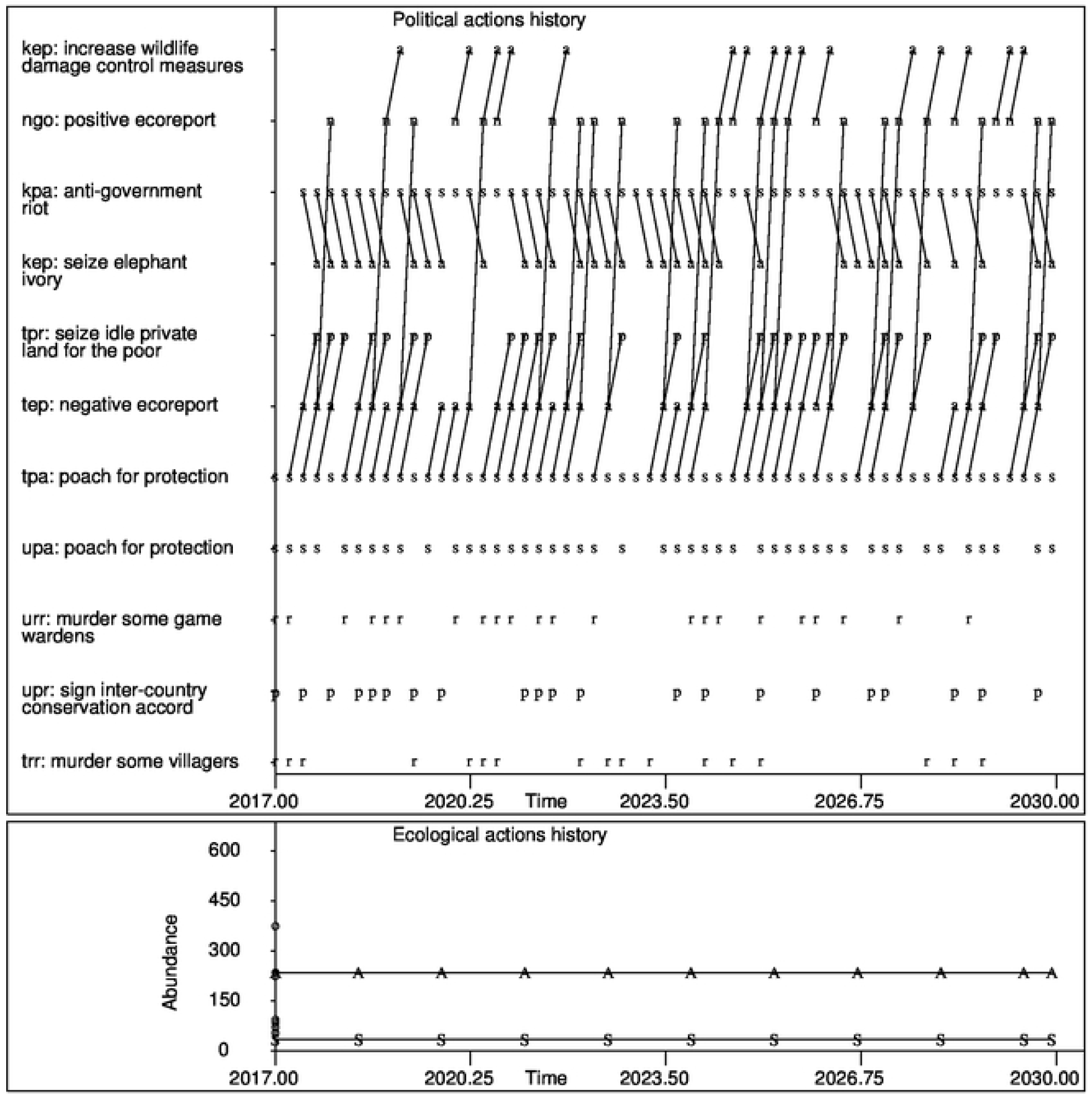
The cheetah EMT simulator’s actions history under the MPEMP. See Figure 2 for symbol legend. Lines connect action-reaction sequences. For example, one frequent action sequence in Tanzania is poaching, followed by a negative ecosystem status report, followed by a land gift to the poor.

This plan’s *ψ* value is 0.845 meaning that this plan is not expected to face severe resistance to its implementation.

## 4 Discussion

A model has been described of the political and ecological processes at play that characterize the dynamics of an ecosystem being impacted by and impacting several different groups of humans. An integrated suite of statistical methods has been presented for assessing its credibility, and computing politically feasible ecosystem management plans with it. Free software has been demonstrated that implements these methods on high performance computing platforms as one cost-effective way to support the lengthy computations that these methods entail. These contributions for the first time, enable ecosystem managers to develop credible models with which to manage an ecosystem that contains endangered species. Given the unprecedented decline in the earth’s biodiversity, the potential impact of this contribution is difficult to overstate.

The EMT procedure given in this article can be used to build political-ecological models for other ecosystem management challenges such as air quality, freshwater pollution, soil contamination, and waste management. But, as indicated by the consistency analysis of the cheetah EMT simulator, current computing resources can support the simultaneous fitting of only a modest fraction of the parameters of a large, policy-relevant simulator.

### 4.1 Other statistical procedures for credibility assessment

The first four simulator jobs described herein do not support in-sample GOF tests nor model selection statistics. A Monte Carlo hypothesis test of a model’s GOF, however, could be found by first building a contingency table whose columns partition the action history’s time interval into 10 or so subintervals, and whose rows index unique out-combinations in the actions history. Each cell in this table holds the observed number of out-combinations in its time subinterval along with the number of out-combinations generated by the model over this time subinterval. The observed chi-squared test statistic for this table would be computed. A Monte Carlo technique would be used to find the p-value for this GOF hypothesis test because there are dependencies across the time subintervals (see [77, pp. 20-22]). This would be done by simulating a large number of Monte Carlo action histories using the estimated model and computing the chi-squared test statistic for each. Finally, the p-value would be found as the fraction of these test statistic values that exceed the observed test statistic value. See [78] for a discussion of this technique.

In agreement with Yarkoni and Westfall [34], however, the present author believes that the final arbitrar of a good model ought to be its out-of-sample prediction error rate. Note that the prediction error rate estimator of Section 2.3.1 is an out-of-sample estimator.

To address the model parsimoniousness goal of model selection procedures, a model of a political-ecological system whose simulator exhibits low prediction error rate, might be made more parsimonious by first setting those parameters whose confidence intervals include zero to very small, fixed values – and then re-computing the model’s prediction error rate to verify that this now more parsimonious model continues to perform at the desired level. See [79] for the reason why confidence intervals may be used to conduct hypothesis tests.

### 4.2 Automatic data streams

As exemplified by the modest cheetah abundance sample size reported in Section 3.1, a limiting factor for applying the suite of statistical methods described herein is the continuous availability of observations on many ecosystem metrics. In other words, to keep a political-ecological simulator relevant for policymaking, the simulator should be regularly refitted to data as new political-ecological is acquired. This regular activity is made more convenient if automatically-acquired streams of political-ecological data are continuously available. See [37] for techniques to create and read such streams.

### 4.3 Funding cluster computer time

HPC providers that offer their compute cycles on the open market include (a) the SDSC [76], (b) Ohio State University’s Supercomputer Center [80], and (c) the private firm, Sabalcore Computing Inc. [81]. Some of these providers allow users to purchase one or more compute nodes for their own, dedicated use. But investing in these so-called “condominium compute nodes” does little to help a user gain access to large numbers of compute nodes.

Until cluster computers become affordable for ecosystem managers, these managers can meet their computing requirements in the face of uneven funding via a JavaSpaces program running on their in-house family of workstations. There is no setup or special software needed other than assigning an IP address to each workstation and installing (free) JAVA and (free) GigaSpaces [66] on each workstation. An important characteristic of this approach is that computing costs are now part of the agency’s office computer budget, i.e., capital expenditures rather than the agency’s budget for services, e.g. consultant fees. As mentioned in Section 2.5, however, a cluster of workstations may not be as reliable nor as fast as a cluster computer.

## 5 Conclusions

The five simulator jobs developed and demonstrated herein show that models of political-ecological systems can be built, statistically estimated, and subjected to rigorous credibility assessment. They can also be used to form ecosystem management policies. But running these jobs can require large amounts of computation. Coding and running them as MTC applications is one way to make them maintainable, financially feasible, and timely. The mathematics and computer code needed to perform such computations have been presented and demonstrated herein. All of this code may be downloaded from [39].

The future of ecosystem management lies in finding workable policies that address not only what needs to be done to conserve ecosystems under anthropogenic pressure, but also the needs and aspirations of those humans who interact with such ecosystems. Building models of these political-ecological systems can help address these challenges but new computational approaches are needed to discover effective and politically implementable management actions from these models. This article provides one such new approach.

## Supporting information captions

**S1 Appendix.** File name: `s1.pdf`. Shell scripts, guidance, and model documentation.

**S2 Data.** File name: `s2.txt`. Observed actions history for the Cheetah EMT simulator.

